# Global Acetylome Profiling Reveals Extensive Lysine Acetylation in *Acinetobacter baumannii* Bacteriophage Proteins

**DOI:** 10.64898/2026.04.27.721073

**Authors:** Rui Chen, Hongyan Zhou, Jianjun Li, Wangxue Chen, Danielle L. Peters

## Abstract

Post-translational modifications (PTMs) are increasingly recognized as potential regulators of bacteriophage–host interactions, yet their functions remain largely undefined. Here, we present the first proteome-wide characterization of lysine acetylation in a bacteriophage, using *Acinetobacter baumannii* myovirus vB_AbaM_DLP3 as a model to investigate host-imposed modification during infection. An optimized proteomic workflow that eliminates chemical artifacts and improves peptide identification accuracy enabled detection of 86 acetylated proteins, representing nearly one-third of the DLP3 proteome. Comparative analysis of purified virions and infected host cells revealed distinct acetylation profiles, consistent with host-driven, transient modifications that are removed or lost after lysis. Functional annotation showed that acetylation predominantly targets proteins involved in DNA replication, genome packaging, and structural assembly. Structural modeling of the major capsid and replication-associated proteins demonstrated that acetylation neutralizes positive surface charges, potentially weakening DNA binding and intercapsomer interactions critical for replication and capsid cohesion. Together, these findings establish lysine acetylation as a widespread and dynamic feature of phage infection and identify host-imposed acetylation as a potential regulatory mechanism that modulates phage replication, assembly, and infectivity. This work expands the molecular framework of phage-host interactions, providing a foundation for future studies aimed at understanding how bacterial PTMs influence phage function and for engineering phages resistant to host-mediated inhibition.

**Importance:** Bacteriophages are being developed as alternatives to antibiotics, yet little is known about how bacterial hosts chemically modify phage proteins during infection. This study reveals that *A. baumannii* imposes extensive lysine acetylation on its infecting phage DLP3, affecting proteins involved in DNA replication and structural assembly. By demonstrating that acetylation neutralizes positive surface charges and is largely absent after phage release, this work identifies host-driven acetylation as a previously unrecognized, reversible mechanism that may influence phage infectivity. Understanding how bacteria use post-translational modifications to interfere with phage function provides new insight into phage-host molecular interactions and informs strategies for engineering therapeutic phages resistant to host-imposed inhibition.

## Introduction

Bacteriophage-based therapy is a promising approach to tackle the global health challenges associated with antimicrobial resistance(1). Recent clinical trials and compassionate-use studies have demonstrated the safety and efficacy of personalized phage therapy, supporting its potential as a complementary or alternative strategy to antibiotics (2, 3). Advances in synthetic biology have enabled rational phage engineering to expand the host range, improve pharmacokinetics, and optimize large-scale production(4).

To effectively engineer therapeutic phages, a fundamental understanding of phage-host interactions at the molecular level is required. During infection, bacteriophages must control the host’s transcriptional and translational machinery while evading bacterial defense systems. This dynamic interplay involves multiple layers of regulation, including post-translational modifications (PTMs)(5), which reversibly alter protein chemistry and function. PTMs such as phosphorylation, acetylation, and ADP-ribosylation are increasingly recognized as key regulators of enzymatic activity, DNA binding, and protein-protein interactions (6).

Although protein acetylation is widespread in bacteria, its role in phage-host interactions remains largely unexplored. To date, only a few isolated studies have reported acetylation events associated with phage infection (7–10), and none have systematically examined the phage acetylome. Interestingly, some phages encode acetyltransferases that acetylate host CRISPR–Cas effectors, thereby disabling bacterial immunity (11). These findings suggest that acetylation could represent a critical regulatory mechanism in both phage offense and host defense; however, the extent and functional consequences of acetylation within phage proteins remain unknown.

Here, we present the first comprehensive proteome-wide analysis of lysine acetylation in a bacteriophage, using *Acinetobacter baumannii* phage vB_AbaM_DLP3 as a model. By optimizing the sample preparation and database search strategies to improve the recovery and accuracy of acetylated peptide identification, we revealed extensive acetylation across the phage proteome and uncovered how host-imposed acetylation may dynamically regulate phage replication and structural assembly. This study defines the landscape of phage acetylation and establishes a foundation for understanding how PTMs shape phage-host molecular warfare.

Deciphering host-imposed acetylation mechanisms could inform the design of engineered phages resistant to bacterial PTM-based defense mechanisms.

## Materials and Methods

All chemicals and reagents were purchased from Sigma-Aldrich (St. Louis, MO, USA) unless specified otherwise. LC-grade water and acetonitrile were purchased from Fisher Scientific (Ottawa, Ontario, Canada) for mass spectrometric analysis.

### Phage and Host Preparation

*Acinetobacter* myovirus DLP3 (vB_AbaM_DLP3; accession PQ034586) was propagated using a capsule mutant of *A. baumannii* strain AB5075 (AB5075^cm^) as previously described(12). Phage lysates were clarified by centrifugation (12,000 x g, 10 min, 4 °C), filtered (0.22 µm PES), purified by CsCl density centrifugation, and buffer-exchanged into 100 mM NaCl, 8 mM MgSO_4_⋅7H_2_O, 50 mM Tris-HCl (pH 7.5). Stocks were titrated by spot plating and stored at 4 °C.

For infection assays, AB5075^cm^ cultures were infected with DLP3 at MOI 1 or 3 and incubated for 15 min at 37 °C with shaking. The cells were pelleted (16,000 × g, 3 min, 4 °C), lysed in 1 % SDS, 100 mM Tris-HCl (pH 7.4) with protease inhibitors, and sonicated (Branson Sonifier® SFX250, 6 cycles, 10 s on/30 s off, 30 % amplitude). Debris was removed by centrifugation (15,000 × g, 5 min, 4 °C).

### Protein Extraction and Digestion Optimization

Proteins were precipitated by adding 10 volumes of cold acetone, incubated overnight at −20 °C, then pelleted and washed three times with acetone. Pellets were resolubilized in 8 M urea, 0.1 % RapiGest (Waters, Milford, MA) or 0.5 % deoxycholate (DOC) in 100 mM Tris-HCl (pH 7.4). Protein concentrations were determined by DC Assay (Bio-Rad) and standardized. Samples were reduced with 10 mM dithiothreitol (56 °C, 45 min). Proteins solubilized in urea and RapiGest were alkylated in the dark with 20 mM iodoacetamide (RT, 25 min). DOC solubilized proteins were alkylated with 20 mM chloroacetamide (RT, 25 min). Proteins were digested overnight with trypsin (sequence grade, Promega, Madison, WI, USA) at a ratio of 1:50 (enzyme:substrate). To remove surfactants, RapiGest and DOC digests were acidified with 10 % trifluoroacetic acid (TFA; 37 °C, 10 min) and centrifuged to collect supernatants. The resulting peptides were desalted using a HyperSep C18 SPE cartridge (Thermo Scientific, Waltham, MA, USA).

### Immunoenrichment of Acetylated Peptides

Acetylated peptides were enriched using procedures described previously (13). Briefly, 200 µg of dried protein digest was resuspended in PTMScan® IAP buffer (Cell Signaling Technology, Danvers, MA, USA) and centrifuged (10,000 × g, 5 min, 4 °C) to remove insoluble pellets.

Acetylated peptides were enriched using PTMScan® acetyl-lysine motif [Ac-K] immunoaffinity beads by incubating the peptide mixture with the beads for 2 hr at 4 °C on a rotator. The beads were collected with a magnet, washed twice with cold IAP washing buffer, then thrice with H_2_O. The peptides were eluted by incubating the beads with 55 μl of 0.15 % (v/v) TFA for 10 min with mixing. The supernatant was collected (2,000 × g, 30 s) and dried by Speed Vac.

### LC-MS/MS Analysis and Data Processing

Samples were analyzed using an Exploris 480 mass spectrometer (Thermo Fisher Scientific, Bremen, Germany) coupled to an Ultimate 3000 UHPLC. Peptides were loaded using an Acclaim PepMap 100 trap column (C18, 5 μm, 100Å, Thermo Scientific, Waltham, MA) onto a Waters BEH C18 column (100 µm × 100 mm) packed with reverse phase beads (1.7 µm, 120Å pore size, Waters, Milford, MA). A 45-min gradient of ACN (5-30 % [v/v] in 0.1 % formic acid [v/v]) was used at an eluent flow rate of 500 nL/min to analyze all the samples. Data-dependent acquisition was performed with a cycle time of 2 s, and a full ms1 scan (resolution: 60, 000; AGC target: standard; maximum IT: 50 ms; scan range: 360-1360 *m/z*) was preceded by subsequent ms2 scans (resolution: 15,000; AGC target: standard; maximum IT: 150 ms; isolation window: 2 *m/z*; scan range: 200-2000 *m/z*; NCE: 30). To minimize repeated sequencing of the same peptide, dynamic exclusion was set to 60 s, and the option of excluding isotopes was activated.

The mass spectrometry proteomics data were deposited in the ProteomeXchange Consortium via the PRIDE (14) partner repository with the dataset identifier PXD076213.

### Database Search, Quantification, and Statistical Analysis

Spectra were searched using MSfragger 4.1 (15) within the FraggerPipe platform using default settings against a composite database (11,587 entries) comprising DLP3 (NCBI, 265 entries), *Acinetobacter baumannii* AB5075 (NCBI, 4566 entries), *Saccharomyces cerevisiae* (strain ATCC 204508 / S288c, Uniprot, 6,735 entries), and the common Repository of Adventitious Proteins (cRAP). A decoy database was generated by reversing target protein sequences and adding them to the target database. Trypsin cleavage at the KR/P was used to determine protease specificity. Both the precursor and fragments tolerance were set at 20 ppm.

Carbamidomethylation of cysteine was used as a fixed modification. N-terminal acetylation and methionine oxidation were set as the variable modification.

To evaluate chemical artifacts, the carbamylation of lysine was included as a variable modification, either with or without isotopic error. Acetylation of lysine was included as a variable modification, and the correction of the isotopic error was disabled. All of the identification results were filtered using an FDR <1 % for PSM, peptides, and proteins. Label-free quantification of the proteome was conducted with Ion Quant (v 1.10) (16) with matches between runs. Statistical analysis was performed using GraphPad Prism 10 using a paired t-test.

### Bioinformatics

Functional annotation was performed using InterProScan (17, 18) and protein BLAST (accessed 2026-02-26) (19, 20). Selected proteins were modelled with and without acetylation using AlphaFold3 (21). Electrostatic surface potentials and structural alignments were analyzed in UCSF ChimeraX (22) v1.11.1 using FoldSeek (23) and Matchmaker (24).

## Results

### Improved Proteome Coverage and Elimination of Carbamylation

To facilitate acetylome analysis, protein extraction with lysis buffer containing SDS was performed to ensure efficient extraction and sterilization of both phages and bacteria. The precipitated proteins were re-dissolved in urea buffer, which is widely used and suggested by the manufacturer for acetylation enrichment using an acetylated lysine antibody(13) (Figure 1A).

**Figure 1.**
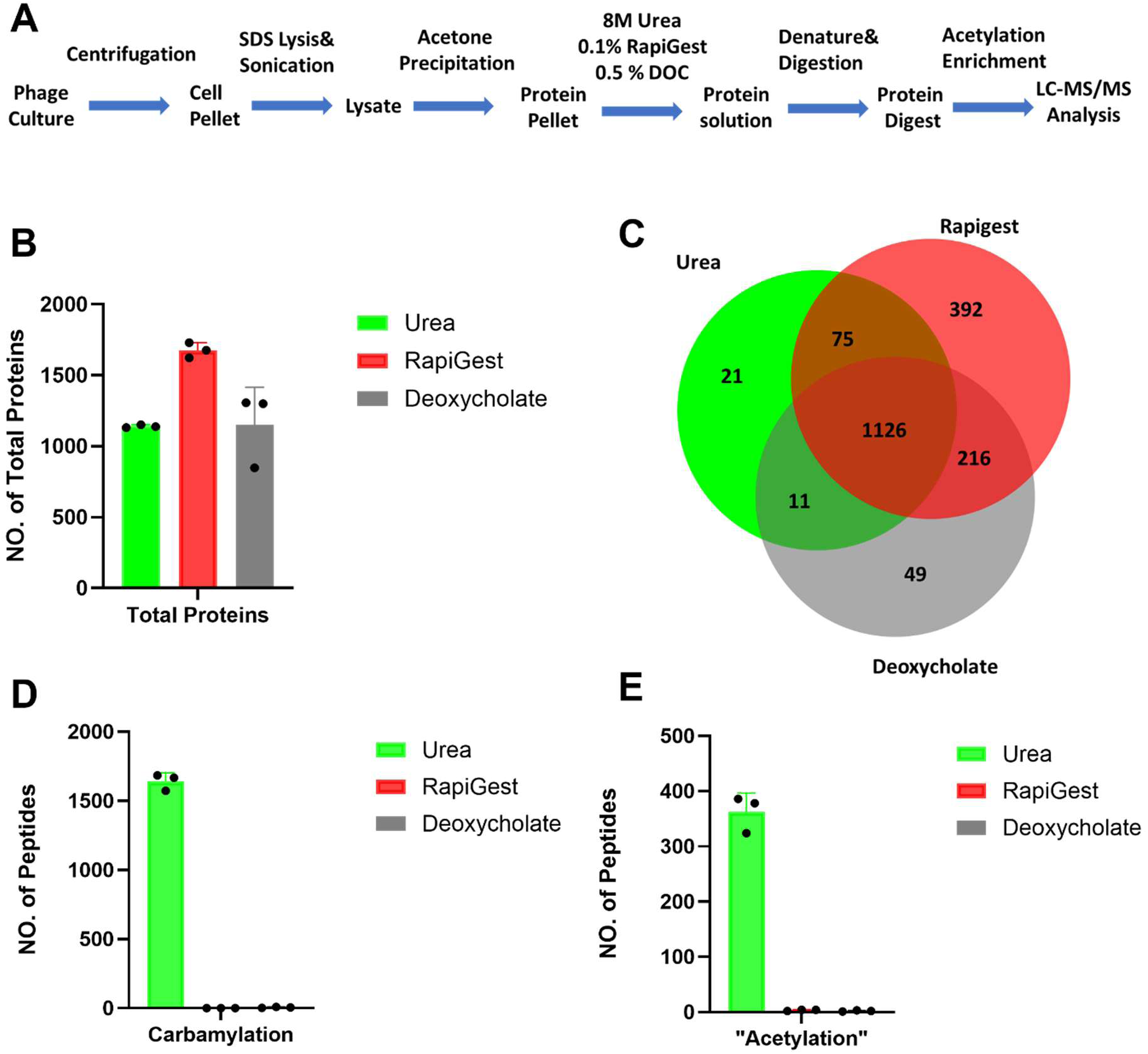
Optimization of sample preparation to increase proteome coverage and reduce chemical artifacts. **A)** Scheme of the workflow for the acetylation analysis from bacteriophage by proteomics; **B)** Comparison of the protein recovery of three different solutions from protein precipitate; **C)** Overlap of proteome coverages from three solutions; **D)** Comparison of carbamylation introduced by three solutions; **E)** Comparison of potentially falsely assigned “acetylation sites” with three solutions.

However, it has been reported that using urea for protein extraction introduces a side reaction of unmodified lysine as urea generates cyanate when heated(25). Carbamylated peptides were co-purified by the antibody as its composition (HC_2_NO, mass=43.0058) was too close to acetylation (H_2_C_2_O, mass=42.0106). Thus, it is necessary to eliminate artifacts while achieving a high proteome coverage. For this purpose, the performance of several protein recovery solutions (urea, deoxycholate and RapiGest) was compared before the enrichment of acetylated peptides (Figure 1B).

Among the three methods tested, RapiGest proved to be the most effective, enabling the identification of the highest number of phage and bacterial proteins at a ratio of 1:1 (MOI=1). The identified proteins also contained a few proteins from *Saccharomyces cerevisiae*, which is unremarkable as yeast extract is a component of the growth media used for propagation. Among the 1841 proteins identified from all three methods, only 32 (1.6 %) and 60 (3.2 %) proteins were detected with urea and deoxycholate, while 392 (21.3 %) proteins were identified using the RapiGest treatment. This suggested that RapiGest provided the highest proteome coverage for phage proteome analysis (Figure 1C).

We further evaluated the procedure for acetylation analysis by adding either carbamylation or acetylation of lysine as a variable modification to the database search of protein digests. The result confirmed that using urea resulted in the highest number of carbamylation sites, whereas only a few sites were detected with either deoxycholate or RapiGest (Figure 1D). With urea, 16.4 % of all identified peptides were carbamylated. When searching with acetylation modification, the number of “acetylated peptides” from urea decreased to 400, and most of the spectra were assigned as carbamylation once carbamylation was set as a variable modification (Figure 1E). Thus, these peptides are most likely carbamylated but could be incorrectly identified as “acetylated.”

From digests processed with deoxycholate or RapiGest, several modified sites were identified, and the abundance of these peptides was much lower than that of their unmodified counterparts. A comparison of the extracted peak area of carbamylated peptides from the urea digestion to the acetylated peptides from the RapiGest digest shows that carbamylation has a much higher stoichiometry than acetylation. This indicated that urea digestion could severely affect acetylation identification using the enrichment strategy. Overall, this evaluation suggests that RapiGest is best suited for acetylation analysis using a bottom-up proteomics approach.

### False Assignment of Acetylation by Allowing Isotopic Error

Replacing urea with RapiGest or deoxycholate for protein denaturation and solubilization can prevent false acetylation assignments arising from carbamylation. Understanding how the database search engine produces these misassignments is also essential for reducing false positives during data analysis. To address this, we analyzed the precursor and MS/MS spectra of carbamylated peptides and determined that excessive isotopic error correction, which is used to increase peptide identifications, was the primary source of misassignment, as illustrated in the following examples.

In the first example, an incorrect monoisotopic peak was selected because of an insufficient number of peaks within the isotopic clusters (Figure 2A). The peak at *m/z* 891.48 was selected for MS/MS, and the MS^2^ spectrum was matched to the peptide ALLSPGIELK#ENTTQSTVVQNATGR with a probability of 0.99. However, the calculated molecular weight of this peak does not match the monoisotopic peak of the “acetylated” peptide; rather, it is more likely the monoisotopic peak of a carbamylated species. As acetylation was set as a variable modification, the search engine permitted a one-Dalton error in monoisotopic peak assignment, allowing the MS/MS spectrum to be incorrectly matched to an acetylated peptide because of the high quality of the spectral fit. By changing the variable modification to carbamylation, the spectrum could be correctly identified as a carbamylated peptide without any isotopic error allowance. Thus, over-correction of isotopic errors can lead to false identification of acetylated peptides, especially from un-enriched samples where modified peptides have a much lower abundance.

**Figure 2.**
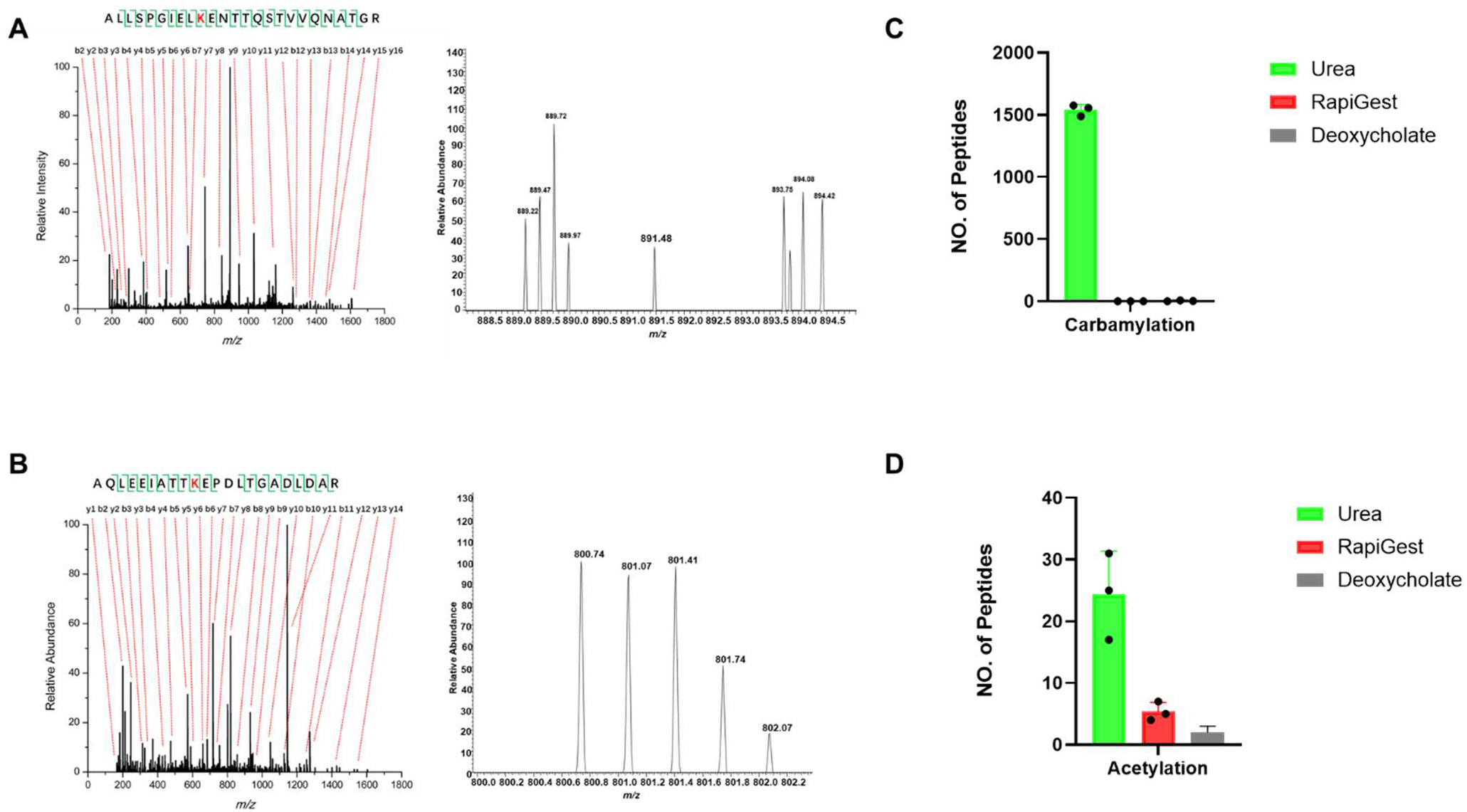
Improving database searching method to further reduce the impact of chemical artifact. **A)** Example of wrongly assigned acetylation by database searching engine due to incorrect selection of monoisotopic peak; **B)** Example of wrongly assigned acetylation by database searching engine due to over correction of the mass of monoisotopic peak; **C)** Number of identified carbamylated peptides remained the same after disabling isotopic error correction; **D)** Number of identified acetylated peptides from total digest decreased significantly after disabling isotopic error correction, proving that mostly falsely assigned acetylation were due to over correction of isotopic error.

The second example resembled the first; however, it was observed even when a complete isotopic cluster was detected. As indicated in Figure 2B, the monoisotopic peak of this peptide should be *m/z* 800.74, which matches the carbamylated form of the peptide AQLEEIATTKEPDLTGADLDAR. However, when acetylation was set as the variable modification, the search engine applied isotopic error correction and generated a non-existent peak at *m/z* 800.40, for which the extracted peak area was zero. When carbamylation was set as a variable modification, the extracted peak area of this peptide increased to 6.5E+8. This finding proves that allowing isotopic error introduces false assignments of carbamylation as acetylation, and is further supported by comparing search results with different isotopic error settings (0/1/2 or 0). When isotopic error correction was not allowed, the number of carbamylation identifications remained unchanged, whereas the number of acetylation dropped dramatically as it almost eliminated all the false identifications (Figure 2C, D). As a result, isotopic error correction was disabled in this study to ensure accurate analysis of phage acetylation.

### Extensive Acetylome Analysis of Phage DLP3

Using improved sample preparation and database search strategies, we used acetyl-lysine antibody enrichment to investigate the acetylation landscape of the DLP3 phage proteome. To maximize coverage of the phage acetylome and understand the role of acetylation during phage-host interactions, both purified phage particles and phage-infected host cells were analyzed.

Among the 265 annotated proteins in the DLP3 proteome, 86 (32.4 %) were found to be acetylated, each containing an average of two modification sites distributed across all structural and functional categories (Supplementary Tables 1 and 2). Importantly, distinct acetylation patterns were observed between the purified phage particles and phage-infected host cells (Figure 3A). Most acetylated phage proteins were detected in the infected host cells, whereas only six acetylated proteins were found in the purified virions. The latter were primarily structural, including the highly abundant major capsid protein (MCP) and low-abundance tail tube protein (Figure 3B).

**Figure 3.**
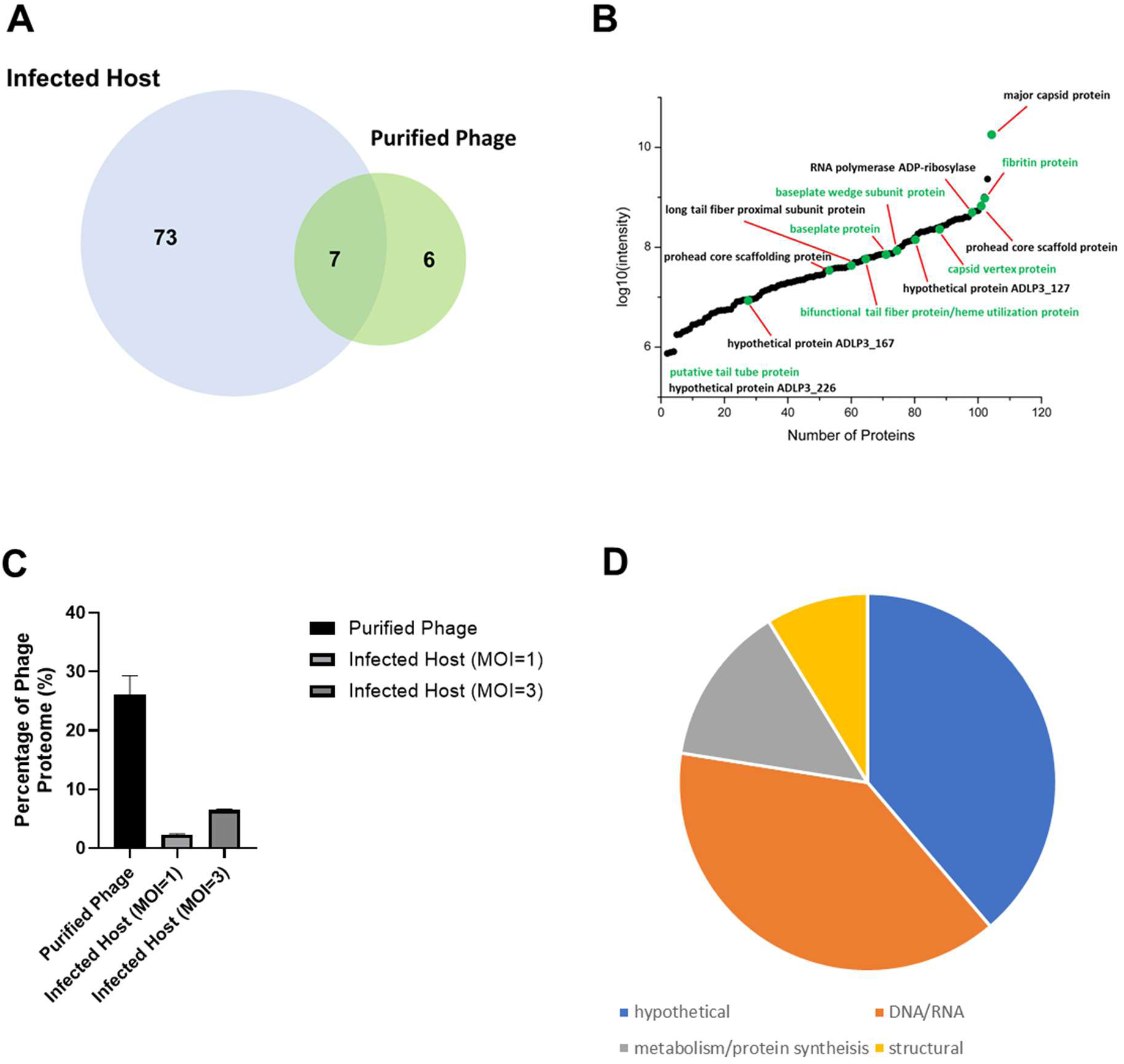
Extensive acetylome analysis of DLP3 phage reveals widespread acetylation in the bacteriophage proteome. **A)** Overlap of acetylated proteins from purified phage and infected host. **B)** Abundance distribution of acetylated phage proteins across the whole proteome. **C)** Percentage of phage proteome from various samples demonstrated that more acetylation was discovered from infected host even though the abundance of phage proteins was lower than that from purified phage. **D)** Functional annotation suggests acetylation is targeted toward proteins involved in DNA/RNA synthesis and replication, and metabolism, during the infection process.

Analysis of protein abundance revealed that acetylation levels were independent of total protein abundance, as supported by a comparison of proteome intensities across samples (Figure 3C). Although phage proteins accounted for > 30 % of the total proteome in purified samples, a higher proportion of acetylated species was observed in infected host cells, suggesting that acetylation may be host-imposed and dynamically regulated during infection (Figure 3D).

Functional enrichment revealed that nearly half of the acetylated proteins were involved in DNA/RNA synthesis, replication, and metabolism, suggesting that host-driven acetylation preferentially targets the proteins required for genome replication and transcriptional control.

### Acetylation of Structural Proteins in Purified Phage Particles

To explore the potential structural consequences, we examined the acetylated proteins identified in purified virions. Notably, several structural components contained multiple acetylation sites, including the long tail fiber proximal subunit (two sites; XEO41410.1), the head outer capsid protein (six sites; XEO41446.1), and the major capsid protein (eight sites; XEO41426.1). In T4-like phages, MCP is encoded by gp23 and is proteolytically processed to gp23* during capsid assembly (26). The essential structural role of gp23* in forming the phage head and mediating intercapsomer contacts prompted a deeper structural analysis of DLP3 MCP.

AlphaFold3 models of both unmodified and acetylated MCP* (residues 69-527) revealed that several acetylated lysines, notably residues K276 and K409, occupy positions analogous to the electrostatic contact zones in T4 gp23*, where complementary charged residues stabilize intercapsomer cohesion (Figure 4A) (27). Electrostatic surface analysis showed that acetylation neutralized these positive surface patches, potentially weakening inter- and intracapsomer interactions (Figure 4B). Additional acetylation sites distributed across the MCP* surface further neutralized the charge, which may further influence capsid packing, stability, or maturation (Figure 4B).

**Figure 4.**
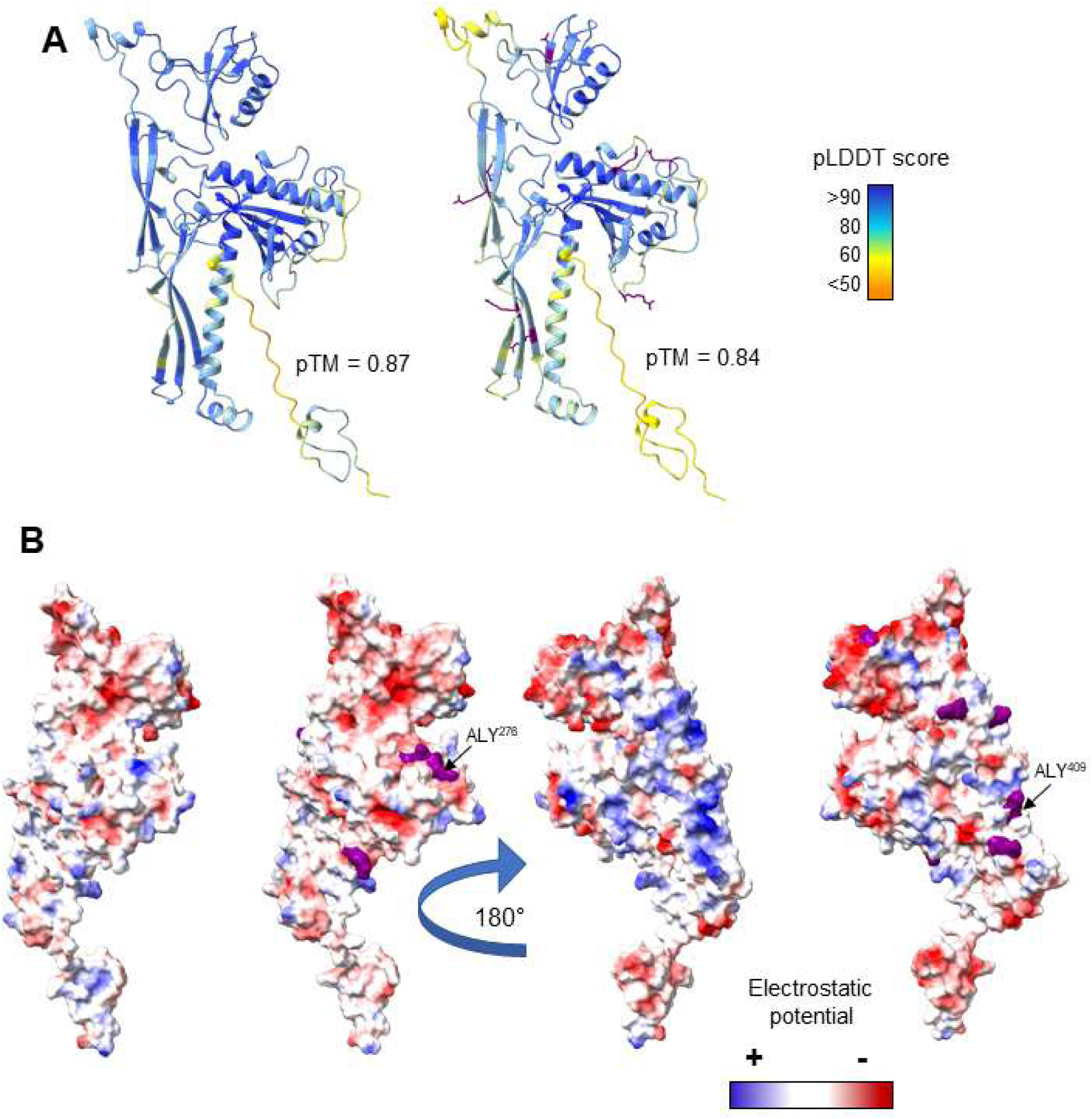
Structural prediction and electrostatic surface analysis of the DLP3 phage major capsid protein (MCP). **A)** AlphaFold3 models of the DLP3 MCP predicted without lysine acetylation (left) and with lysine acetylation (right), colored according to plDDT confidence scores. Predicted pTM scores are shown for each model. Lysine acetylation sites identified by LC-MS/MS are highlighted in purple. **B)** Coulombic electrostatic surface potential maps of the MCP shown in two orientations separated by a 180° rotation. The first and third structures correspond to the unmodified MCP, and the second and forth correspond to the acetylated MCP. Acetylated lysines are highlighted in purple. Acetylation results in localized changes in surface electrostatic potential that may alter MCP surface interactions relevant to capsid assembly or maturation.

Acetylation of MCP was previously detected in marine bacteriophages, and it was suggested that these modifications do not stabilize the capsid architecture; instead, they may play more complex roles in other biological processes (10). Together with our findings on *Acinetobacter* phage DLP3, we suggest that acetylation of MCP and other structural proteins is a host-imposed anti-phage strategy used to modulate capsid assembly or post-assembly stability by disrupting electrostatic interactions critical for virion architecture.

### Acetylation of Replication-Associated Proteins in Infected Host Cells

Most acetylated phage proteins are modified within host cells and lost after lysis, suggesting that acetylation is transient and reversible. Functional annotation identified several replication- and transcription-associated proteins with extensive acetylation, including the medium and large DNA Topoisomerase II (XEO41419.1, XEO41435.1) proteins, DNA Polymerase I (XEO41447.1), and the hypothetical protein ADLP3_254 (XEO41462.1), suggesting that acetylation could be imposed by the host to hinder transcription and translation.

ADLP3_254 exhibited the highest acetylation density, with 11 acetylated lysine sites. InterProScan analysis identified strong homology to the single-stranded DNA-binding (ssDNA-binding) protein Gp32 of *Escherichia* phage T4 (IPR046395), which is essential for DNA replication, recombination, and repair (28) (29). Structural modeling of ADLP3_254 using AlphaFold3 produced high-confidence predictions for residues 29-239, which corresponded to a well-defined core region (Figure 5A). The overall predicted template modeling (pTM) scores were 0.77 for the acetylated model and 0.70 for the unmodified model, suggesting slightly greater structural confidence when the acetylated lysine PTMs are included.

**Figure 5.**
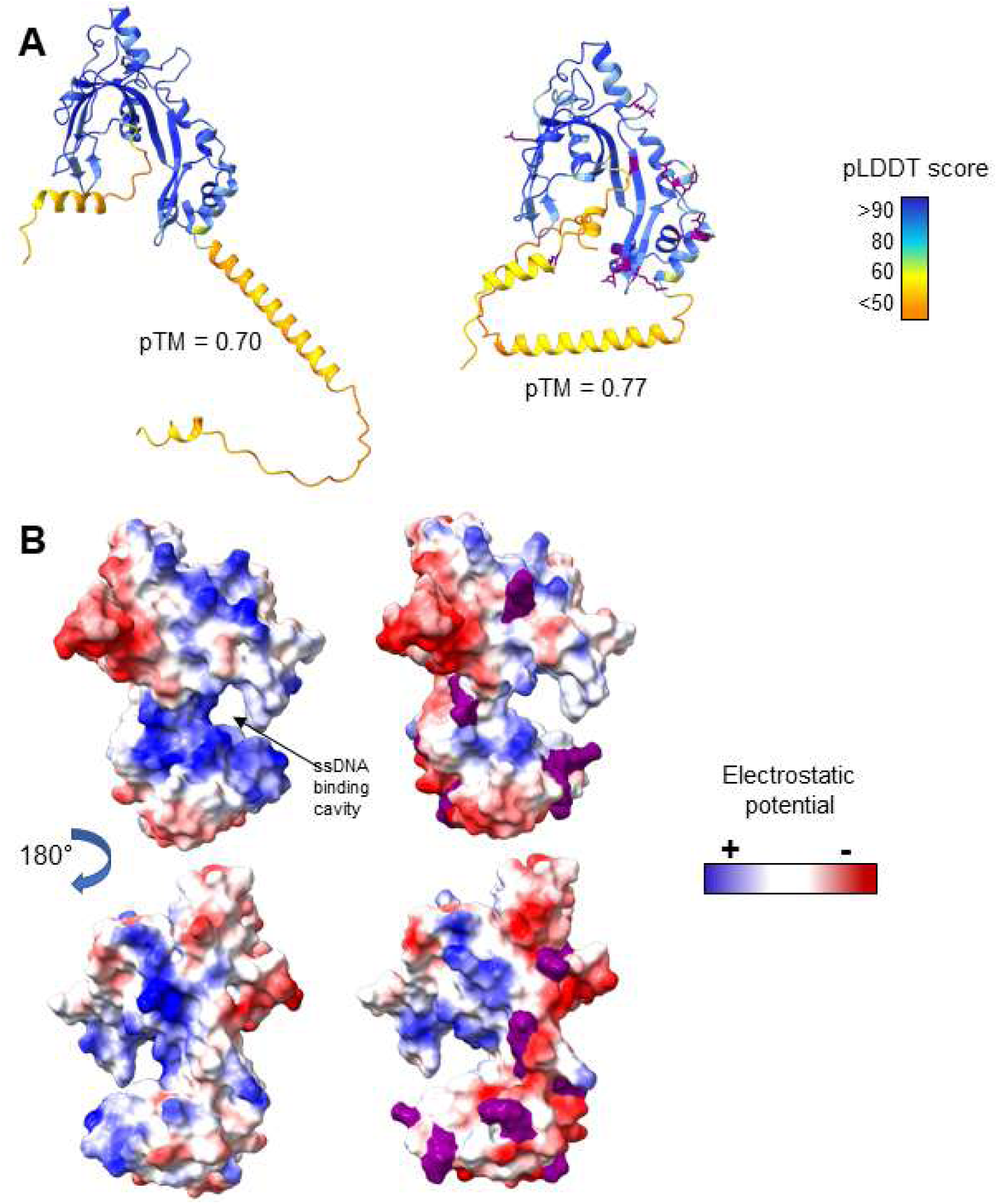
Structural prediction of the full-length ssDNA binding protein ADLP3_254 and electrostatic effects of lysine acetylation. **A)** AlphaFold3 models of full-length ADLP3_254 predicted without lysine acetylation (left) and with lysine acetylation (right), colored according to plDDT confidence scores. Predicted pTM scores are indicated for each model. Both models show high confidence across residues 29-239, corresponding to the structured core region, with lower confidence in the N- and C-terminal regions. Acetylated lysine residues are highlighted in purple. **B)** Coulombic electrostatic surface potential maps of the structured core region of ADLP3_254 in the absence (left) and presence (right) of lysine acetylation, shown in two orientations separated by a 180° rotation. Acetylated lysines are highlighted in purple.

Structural homologues of both ADLP3_254 models were identified using FoldSeek against the Protein Data Bank (PDB). The closest matches were the ssDNA-binding proteins from *Escherichia* phages RB69 (PDB ID: 2A1K) and T4 (PDB ID:1GPC) (Supplementary Table 4). Structural alignment in UCSF ChimeraX using the Matchmaker tool revealed a strong similarity between ADLP3_254 and these homologues. The unmodified ADLP3_254 model aligned with the RB69 ssDNA-binding protein with a root mean square deviation (RMSD) of 0.538 Å across all residues, whereas the acetylated model showed a slightly higher RMSD of 0.647 Å, which is consistent with subtle conformational adjustments associated with lysine acetylation.

To better visualize these effects within the DNA-binding core, the ADLP3_254 model was truncated to include residues 29-239 and remodeled with and without lysine acetylation.

Electrostatic surface-potential analysis of this core region revealed that acetylation neutralized the positive charge within the ssDNA-binding pocket, particularly around lysine residues K224 and K230 (Figure 5B). This localized charge reduction is predicted to weaken the electrostatic interactions with the negatively charged DNA backbone, providing a plausible mechanism by which host-imposed acetylation may impede DNA replication and repair.

Acetylation results in localized changes in surface electrostatic potential, including in the predicted ssDNA-binding cavity, and may influence nucleic acid binding.

The ADLP3_254 protein also served as an example to demonstrate the accuracy of acetylation identification. Two adjacent acetylation sites, K224 and K230 (Figure 6A, B, and C), were detected in different peptides, each showing high-quality fragment ion matches that precisely pinpointed modification sites. The unmodified peptide, DKFNSYEK, was also identified. A comparison of its matched spectrum with that of the acetylated peptide revealed a mass difference of 42 Da between a series of b ions, clearly confirming the presence of acetylation. Further, the retention time differences are also consistent with acetylation, as the unmodified peptide was eluted at 12.6 min, while the acetylated peptide was eluted at 19.7 min due to the increased hydrophobicity added by the acetyl group.

**Figure 6.**
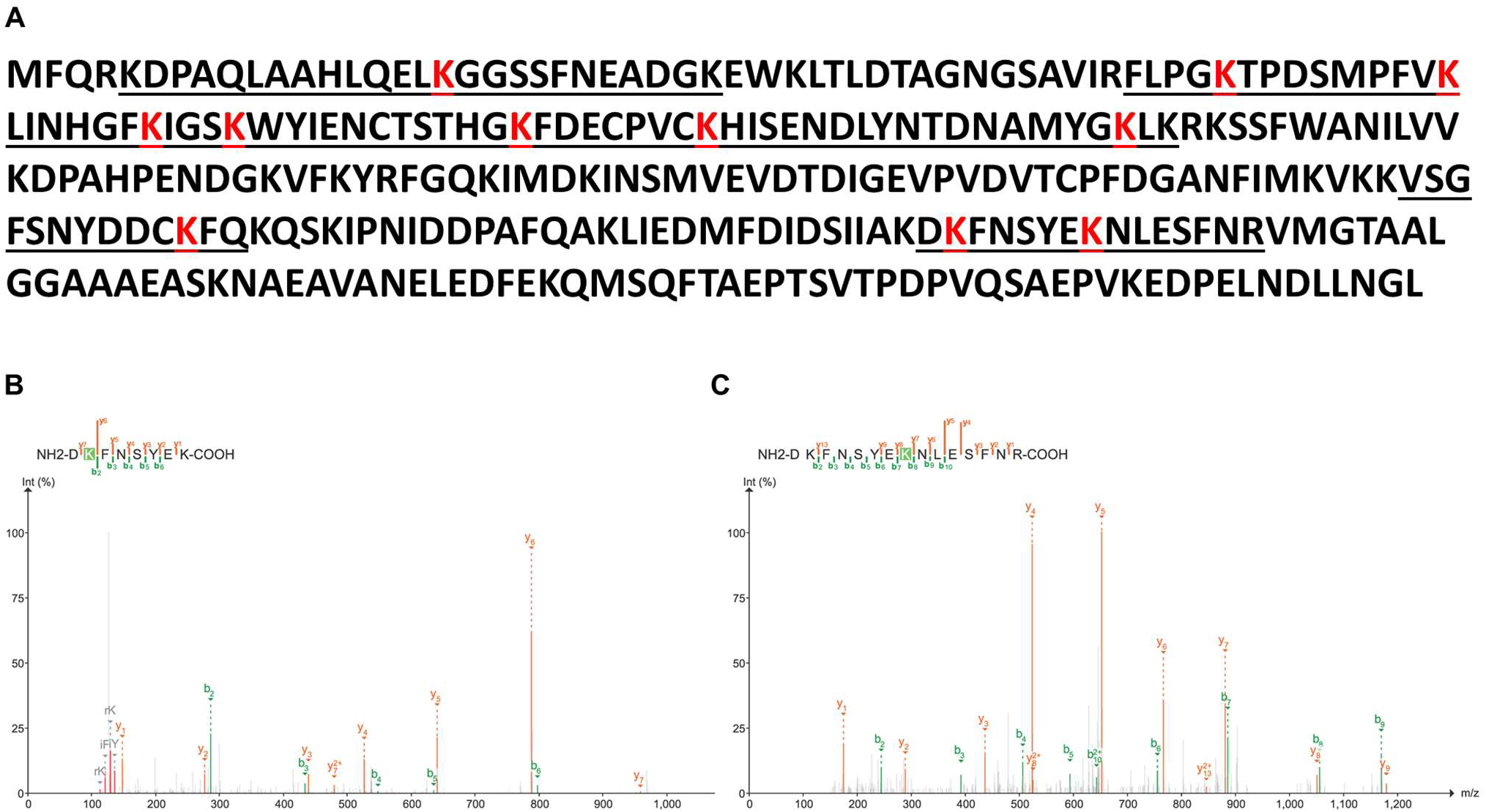
Extensive and accurate identification of acetylation in phage proteins exemplified by ADLP3_254. **A)** protein sequence of ADLP3 with all the acetylated peptides (underlined) and lysine site (in red) labeled; **B)** Matched MS/MS spectrum of acetylation on K224 and sequence of identified peptide; **C)** Matched MS/MS spectrum of acetylation on K230 and the sequence of identified peptide. The two adjacent sites were identified by different peptides and varied with stringent manual validation.

Taken together, our results revealed that lysine acetylation is a widespread and multifaceted modification of the DLP3 phage proteome. Structural modeling of both virion and host-associated proteins demonstrated that acetylation neutralizes positive surface charges, altering electrostatic interactions essential for DNA binding, genome packaging, and capsid cohesion. The prevalence of acetylation in replication and structural proteins within infected host cells, in contrast to its near absence in purified virions, suggests that this modification is transient, host imposed, and functionally targeted. Collectively, these findings define the first detailed phage acetylome and provide mechanistic insight into how host driven PTMs may regulate viral replication and assembly.

## Discussion

Accurate characterization of post-translational modifications (PTMs) in bacteriophages has been limited by technical challenges associated with complex host backgrounds and low modification abundance. In most proteomic studies of phages, infected host cells are lysed with SDS-PAGE loading buffer followed by in-gel digestion (7, 8, 30, 31). Although this approach provides broad proteomic coverage, PTMs can be lost during in-gel digestion. Modern proteomics workflows favor in-solution digestion, filter aided sample preparation, and single-pot solid-phase-enhanced sample preparation for PTM analysis (32). Accordingly, we used SDS-based cell lysis, acetone precipitation, and in-solution digestion with RapiGest, which yielded excellent recovery of acetylated peptides from both the phage and host samples.

As acetylome analysis of bacteriophages during infection has not been previously reported, the performance of this optimized workflow was compared to that of bacterial acetylome studies. A total of 1,388 acetylated peptides from 545 proteins were identified in *A. baumannii* strain AB5075 (Supplementary Table 3), which is substantially higher than those previously reported for other *A. baumannii* strains(33, 34). Disabling isotopic error correction during database searching further improves accuracy by eliminating false assignments arising from chemical artifacts, thereby increasing confidence in both host- and phage-derived acetylation sites.

Despite these advances, the quantitative comparison of acetylation levels between samples remains difficult because the host proteome background varies considerably. Thus, the functional roles of acetylation are best inferred from the location and biological functions of the modified proteins. As shown in histone studies, acetylation neutralizes the positive charge of lysine residues, weakening the electrostatic interactions between negatively charged DNA and positively charged histones(35). Analogously, the host-mediated acetylation of phage proteins may reduce DNA-binding affinity, thereby impeding phage genome replication. The predominance of acetylated phage proteins involved in DNA/RNA synthesis and metabolism supports this hypothesis, suggesting a host-imposed defence mechanism that modulates phage replication through charge neutralization.

Furthermore, the marked reduction of acetylated phage proteins in purified virions compared with that in infected host cells indicates that acetylation is reversible and largely absent after phage release. This suggests that acetylation is imposed by the host and subsequently removed, possibly by the host or phage deacetylases, once the assembly is complete. Investigating these enzymatic mediators will clarify whether acetylation is lost during lysis or persists transiently in newly formed virions. The observation that acetylation targets conserved lysine residues in T4-like myoviruses (RB69 and T4) implies an evolutionarily constrained mechanism through which hosts exploit PTMs to interfere with viral functions.

Acetylation was also detected in structural proteins involved in host recognition, binding and genome delivery, including several tail and baseplate components. For example, the baseplate wedge subunit ADLP3_178 was found to be acetylated; its counterpart in phage T4 interacts with the host cell receptor, regulates tail sheath contraction in myoviruses, and initiates genome ejection (36). The presence of acetylation of analogous structural proteins in DLP3 suggests that this modification could influence the conformational transitions required for adsorption and infection. In addition to ADLP3_178, several other tail and baseplate proteins contain acetylation sites, indicating that host-imposed acetylation may modulate receptor binding and mechanical activation during infection. These structural modifications could serve as potential targets for rational phage engineering to overcome host defenses, for example, by substituting lysine residues with other amino acids that are less susceptible to acetylation (37).

Overall, our results demonstrated that lysine acetylation is widespread across the DLP3 phage proteome and occurs predominantly within the host environment. Although our structural modeling and electrostatic analyses suggest that acetylation may influence DNA binding, genome packaging, and capsid cohesion through charge neutralization, these effects remain predictive and require direct experimental validation. The observed enrichment of acetylation on replication and structural proteins supports the possibility that host-imposed acetylation modulates phage function, but additional biochemical and genetic studies are needed to confirm these roles and to identify the specific bacterial enzymes responsible. Nevertheless, our findings establish a foundational dataset and analytical framework for investigating post translational regulation in phages. By defining where and how acetylation occurs, this study sets a stage for future functional work to elucidate how host-driven chemical modifications shape phage replication, assembly, and evolution.

## Conclusion

This study provides the first comprehensive characterization of lysine acetylation in a bacteriophage and its host during infection, revealing that more than one-third of the DLP3 proteome is subject to host-imposed modifications. By integrating optimized proteomic workflows with structural modeling, we mapped the distribution of acetylation sites and predicted how this modification neutralizes the positive surface charges on proteins essential for DNA replication and capsid assembly. These findings establish lysine acetylation as a widespread and dynamic feature of phage biology, and suggest that bacteria may exploit this process as part of an antiviral defense strategy.

Although structural modeling indicates that acetylation could alter electrostatic interactions critical to phage function, these effects remain predictive and require direct experimental validation. Nonetheless, our work defines the landscape of phage acetylation and provides a foundation for future investigations into how host driven chemical modifications influence viral replication and assembly. Beyond mapping the phage acetylome, this study underscores the importance of considering post translational regulation as an integral component of phage host dynamics. Future studies should identify the bacterial enzymes mediating acetylation and deacetylation and test whether targeted mutagenesis of acetylation prone residues can yield engineered phages resistant to host-imposed inhibition. Together, these insights open new avenues for exploring PTM-based regulation in viruses and developing next generation therapeutic phages optimized for clinical use.

## Supporting Information

Data are available at ProteomeXchange with the identifier PXD076213, and the DLP3 genome can be accessed through NCBI: PQ034586. The protein models are available upon request.

## Appendixes

**A1.** Supplementary Table 1: list of Identified acetylated peptides from purified DLP3 phage

**A2.** Supplementary Table 2: List of Identified acetylated peptides in DLP3 phage proteins from infected *Acinetobacter baumannii* AB5075

**A3.** Supplementary Table 3: List of Identified acetylated peptides from infected *Acinetobacter baumannii* AB5075

**A4.** Supplementary Table 4: Foldseek and Matchmaker data for ADLP3_254, MCP, and Topoisomerase M and L with and without acetylation PTM

## Author Contribution Statement

Conceptualization: RC and DLP; Methodology: RC and DLP; Validation: RC, JL, HZ, WC, and DLP; Formal analysis: RC and DLP; Investigation: RC, HZ, and DLP; Resources: DLP; Data curation: RC, DLP, and HZ; Writing – original draft: RC, DLP, and HZ; Writing – review and editing: DLP, RC, JL, WC; Visualization: RC and DLP; Project administration: DLP; Funding acquisition: DLP.

## Acknowledgements

The authors thank the Human Health Therapeutics Research Center of the National Research Council of Canada for internal funding to conduct this research.

## References

1. Hatfull GF, Dedrick RM, Schooley RT. Phage therapy for antibiotic-resistant bacterial infections. Annual Review of Medicine. 2022;73(1):197–211.

2. Pirnay JP, Djebara S, Steurs G, Griselain J, Cochez C, De Soir S, et al. Personalized bacteriophage therapy outcomes for 100 consecutive cases: a multicentre, multinational, retrospective observational study. Nat Microbiol. 2024;9(6):1434–53.

3. Kim P, Sanchez AM, Penke TJR, Tuson HH, Kime JC, McKee RW, et al. Safety, pharmacokinetics, and pharmacodynamics of LBP-EC01, a CRISPR-Cas3-enhanced bacteriophage cocktail, in uncomplicated urinary tract infections due to Escherichia coli (ELIMINATE): the randomised, open-label, first part of a two-part phase 2 trial. Lancet Infect Dis. 2024;24(12):1319–32.

4. Jia H-J, Jia P-P, Yin S, Bu L-K, Yang G, Pei D-S. Engineering bacteriophages for enhanced host range and efficacy: Insights from bacteriophage-bacteria interactions. Frontiers in Microbiology. 2023;14:1172635.

5. Longin H, Broeckaert N, van Noort V, Lavigne R, Hendrix H. Posttranslational modifications in bacteria during phage infection. Current Opinion in Microbiology. 2024;77:102425.

6. Carabetta VJ, Hardouin J. Editorial: Bacterial Post-translational Modifications. Front Microbiol. 2022;13:874602.

7. Wright BW, Logel DY, Mirzai M, Pascovici D, Molloy MP, Jaschke PR. Proteomic and transcriptomic analysis of microviridae φX174 infection reveals broad upregulation of host Escherichia coli membrane damage and heat shock Responses. MSystems. 2021;6(3):10.1128/msystems.00046-21.

8. Bleriot I, Blasco L, Pacios O, Fernández-García L, López M, Ortiz-Cartagena C, et al. Proteomic study of the interactions between phages and the bacterial host Klebsiella pneumoniae. Microbiology Spectrum. 2023;11(2):e03974–22.

9. Wolfram-Schauerte M, Pozhydaieva N, Viering M, Glatter T, Höfer K. Integrated omics reveal time-resolved insights into T4 phage infection of E. coli on proteome and transcriptome levels. Viruses. 2022;14(11):2502.

10. Wei S, Wang A, Cai L, Ma R, Lu L, Li J, et al. Proteomic Analysis of Marine Bacteriophages: Structural Conservation, Post - Translational Modifications, and Phage – Host Interactions. Environmental Microbiology. 2025;27(4):e70099.

11. Dong L, Guan X, Li N, Zhang F, Zhu Y, Ren K, et al. An anti-CRISPR protein disables type V Cas12a by acetylation. Nature structural & molecular biology. 2019;26(4):308–14.

12. Peters DL, Davis CM, Harris G, Zhou H, Rather PN, Hrapovic S, et al. Characterization of virulent T4-like Acinetobacter baumannii bacteriophages DLP1 and DLP2. Viruses. 2023;15(3):739.

13. Zhang X, Ning Z, Mayne J, Yang Y, Deeke SA, Walker K, et al. Widespread protein lysine acetylation in gut microbiome and its alterations in patients with Crohn’s disease. Nature communications. 2020;11(1):4120.

14. Perez-Riverol Y, Bandla C, Kundu DJ, Kamatchinathan S, Bai J, Hewapathirana S, et al. The PRIDE database at 20 years: 2025 update. Nucleic Acids Res. 2025;53(D1):D543–D53.

15. Kong AT, Leprevost FV, Avtonomov DM, Mellacheruvu D, Nesvizhskii AI. MSFragger: ultrafast and comprehensive peptide identification in mass spectrometry–based proteomics. Nature methods. 2017;14(5):513–20.

16. Yu F, Haynes SE, Teo GC, Avtonomov DM, Polasky DA, Nesvizhskii AI. Fast quantitative analysis of timsTOF PASEF data with MSFragger and IonQuant. Molecular & Cellular Proteomics. 2020;19(9):1575–85.

17. Blum M, Andreeva A, Florentino LC, Chuguransky SR, Grego T, Hobbs E, et al. InterPro: the protein sequence classification resource in 2025<? mode longmeta?>. Nucleic acids research. 2025;53(D1):D444–D56.

18. Jones P, Binns D, Chang H-Y, Fraser M, Li W, McAnulla C, et al. InterProScan 5: genome-scale protein function classification. Bioinformatics. 2014;30(9):1236–40.

19. Altschul SF, Madden TL, Schaffer AA, Zhang J, Zhang Z, Miller W, et al. Gapped BLAST and PSI-BLAST: a new generation of protein database search programs. Nucleic Acids Res. 1997;25(17):3389–402.

20. Altschul SF, Wootton JC, Gertz EM, Agarwala R, Morgulis A, Schaffer AA, et al. Protein database searches using compositionally adjusted substitution matrices. FEBS J. 2005;272(20):5101–9.

21. Abramson J, Adler J, Dunger J, Evans R, Green T, Pritzel A, et al. Accurate structure prediction of biomolecular interactions with AlphaFold 3. Nature. 2024;630(8016):493–500.

22. Meng EC, Goddard TD, Pettersen EF, Couch GS, Pearson ZJ, Morris JH, et al. UCSF ChimeraX: Tools for structure building and analysis. Protein Sci. 2023;32(11):e4792.

23. van Kempen M, Kim SS, Tumescheit C, Mirdita M, Lee J, Gilchrist CLM, et al. Fast and accurate protein structure search with Foldseek. Nat Biotechnol. 2024;42(2):243–6.

24. Meng EC, Pettersen EF, Couch GS, Huang CC, Ferrin TE. Tools for integrated sequence-structure analysis with UCSF Chimera. BMC Bioinformatics. 2006;7:339.

25. Martinez-Val A, Garcia F, Ximénez-Embún P, Martinez Teresa-Calleja A, Ibarz N, Ruppen I, et al. Urea artifacts interfere with immuno-purification of lysine acetylation. Journal of proteome research. 2017;16(2):1061–8.

26. Leiman PG, Kanamaru S, Mesyanzhinov VV, Arisaka F, Rossmann MG. Structure and morphogenesis of bacteriophage T4. Cell Mol Life Sci. 2003;60(11):2356–70.

27. Chen Z, Sun L, Zhang Z, Fokine A, Padilla-Sanchez V, Hanein D, et al. Cryo-EM structure of the bacteriophage T4 isometric head at 3.3-A resolution and its relevance to the assembly of icosahedral viruses. Proc Natl Acad Sci U S A. 2017;114(39):E8184–E93.

28. Cashen BA, Morse M, Rouzina I, Karpel RL, Williams MC. Dynamic structure of T4 gene 32 protein filaments facilitates rapid noncooperative protein dissociation. Nucleic acids research. 2023;51(16):8587–605.

29. Albrecht CS, Israels B, Maurer J, H von Hippel P, Marcus AH. Functional integration of the bacteriophage T4 DNA replication complex: The multiple roles of the ssDNA binding protein (gp32). Biochemistry. 2025;65(2):207–21.

30. Lemay M-L, Otto A, Maaß S, Plate K, Becher D, Moineau S. Investigating Lactococcus lactis MG1363 response to phage p2 infection at the proteome level*[S]. Molecular & Cellular Proteomics. 2019;18(4):704–14.

31. Emslander Q, Vogele K, Braun P, Stender J, Willy C, Joppich M, et al. Cell-free production of personalized therapeutic phages targeting multidrug-resistant bacteria. Cell chemical biology. 2022;29(9):1434–45. e7.

32. Varnavides G, Madern M, Anrather D, Hartl N, Reiter W, Hartl M. In search of a universal method: a comparative survey of bottom-up proteomics sample preparation methods. Journal of proteome research. 2022;21(10):2397–411.

33. Kentache T, Jouenne T, Dé E, Hardouin J. Proteomic characterization of Nα-and Nε-acetylation in Acinetobacter baumannii. Journal of Proteomics. 2016;144:148–58.

34. Liao J-H, Tsai C-H, Patel SG, Yang J-T, Tu I-F, Lo Cicero M, et al. Acetylome of Acinetobacter baumannii SK17 reveals a highly-conserved modification of histone-like protein HU. Frontiers in Molecular Biosciences. 2017;4:77.

35. Sterner DE, Berger SL. Acetylation of histones and transcription-related factors. Microbiology and molecular biology reviews. 2000;64(2):435–59.

36. Taylor NM, Prokhorov NS, Guerrero-Ferreira RC, Shneider MM, Browning C, Goldie KN, et al. Structure of the T4 baseplate and its function in triggering sheath contraction. Nature. 2016;533(7603):346–52.

37. Peng H, Chen IA, Qimron U. Engineering phages to fight multidrug-resistant bacteria. Chemical Reviews. 2024;125(2):933–71.

